# A decade of epigenetic change in aging twins: genetic and environmental contributions to longitudinal DNA methylation

**DOI:** 10.1101/778555

**Authors:** Chandra A. Reynolds, Qihua Tan, Elizabeth Munoz, Juulia Jylhävä, Jacob Hjelmborg, Lene Christiansen, Sara Hägg, Nancy L. Pedersen

## Abstract

**Background:** Epigenetic changes may result from the interplay of environmental exposures and genetic influences and contribute to differences in age-related disease, disability and mortality risk. However, the etiologies contributing to stability and change in DNA methylation have rarely been examined longitudinally.

**Methods:** We considered DNA methylation in whole blood leukocyte DNA across a 10-year span in two samples of same-sex aging twins: (a) Swedish Adoption Twin Study of Aging (SATSA; N = 53 pairs, 53% female; 62.9 and 72.5 years, SD=7.2 years); (b) Longitudinal Study of Aging Danish Twins (LSADT; N = 43 pairs, 72% female, 76.2 and 86.1 years, SD=1.8 years). Joint biometrical analyses were conducted on 358,836 methylation probes in common. Bivariate twin models were fitted, adjusting for age, sex and country.

**Results:** Overall, results suggest genetic contributions to DNA methylation across 358,836 sites tended to be small and lessen across 10 years (broad heritability *M*=23.8% and 18.0%) but contributed to stability across time while person-specific factors explained emergent influences across the decade. Aging-specific sites identified from prior EWAS and methylation age clocks were more heritable than background sites. The 5,037 sites that showed the greatest heritable/familial-environmental influences (*p*<1E-07) were enriched for immune and inflammation pathways while 2,020 low stability sites showed enrichment in stress-related pathways.

**Conclusions:** Across time, stability in methylation is primarily due to genetic contributions, while novel experiences and exposures contribute to methylation differences. Elevated genetic contributions at age-related methylation sites suggest that adaptions to aging and senescence may be differentially impacted by genetic background.

## Introduction

The functional profiles of genes are not static and vary across time, and indeed across the lifespan, in part as a result of different environmental exposures and contexts (M. J. Jones, Goodman, & Kobor, 2015; Lappe & Landecker, 2015; McClearn, 2006; van Dongen et al., 2016). Measurable gene-environment dynamics for behavioral traits are possible due to advances in biotechniques for global epigenetic profiling at, e.g. specific methylation sites in the human genome. Epigenetic changes may be critical to the development of complex diseases, accelerated aging, or steeper declines in cognitive and physical functioning with age (Lappe & Landecker, 2015). Understanding epigenetic changes over time in the elderly may identify pathways of decline or plasticity (e.g., maintenance or even boosts in functioning) during the aging process and help with elucidating the biology of aging and survival.

Epigenetic modifications resulting in altered gene expression may occur due to a number of processes, including direct methylation of DNA (P. A. Jones & Takai, 2001). DNA methylation results from intrinsic-programmed factors as well as nongenetic processes that may arise due to prenatal or early life exposures or at later points in development (Gottesman & Hanson, 2005; Kanherkar, Bhatia-Dey, & Csoka, 2014; Torano et al., 2016). DNA methylation is characteristically produced by the addition of a methyl group to the DNA molecule cytosine within cytosine-guanine dinucleotides (CpGs), at an estimated 28 million sites across the human genome (Lovkvist, Dodd, Sneppen, & Haerter, 2016). Dense regions of CpGs referred to as ‘islands’ and represent about 5% of CpGs occurring in the genome (about 20,000 total) and often reside in promotor regions (Vinson & Chatterjee, 2012); in addition, surrounding ‘shores’ and ‘shelves’ to these islands are of interest and may be differentially methylated compared to islands (M. J. Jones et al., 2015). The addition of methylation tags to CpG sites is associated with altered gene expression, typically by interfering with or silencing gene transcription although upregulation of gene expression has been documented (C. Wang et al., 2019), and may differentially occur in cells across multiple tissue types including brain, muscle and leukocytes (Fernandez et al., 2012). Methylation tags can be removed as a consequence of exposures as well, leading to dynamics in expression across time (Kanherkar et al., 2014).

Although epigenetic variation is largely attributed to environmental factors (Hannon et al., 2018; Torano et al., 2016; van Dongen et al., 2016), there is evidence for genetic contributions to variation in methylation across the epigenome (Hannon et al., 2018; Torano et al., 2016; van Dongen et al., 2016). Average heritabilities of 16.5 – 19.0% have been reported across sites in the Illumina 450k chip array from whole blood and common environmental influences of 3.0% to 12.6% (Hannon et al., 2018; van Dongen et al., 2016). Stronger evidence of common environment has been reported in young adulthood (18 years) at 12.6% (after correction for cell types; Hannon et al., 2018). Moreover, cross-sectional work suggests that there may be smaller heritable components by mid-adulthood (18%) than young adulthood (21%) (van Dongen et al., 2016),

Epigenetic changes may accelerate over time, whereby changes in gene expression due to exposures become more abundant and salient to phenotypic changes, hence potentiating the development of health and aging conditions earlier in life. Indeed, methylation is correlated with age (Ciccarone, Tagliatesta, Caiafa, & Zampieri, 2018; van Dongen et al., 2016), is used to define biological clocks that may more closely track biological aging (Field et al., 2018), and is associated with mortality (Y. Zhang et al., 2017) and a number of physical and neuropsychiatric health traits (Kanherkar et al., 2014; Lappe & Landecker, 2015). Longitudinal studies of twins represent a valuable approach to evaluate genetic and environmental contributions to stability and change in methylation across the methylome (Tan, Christiansen, von Bornemann Hjelmborg, & Christensen, 2015). Investigations of etiological contributions have relied primarily on crosssectional data (Hannon et al., 2018; van Dongen et al., 2016) and have addressed age-related differences (van Dongen et al., 2016) but not change. We evaluate individual differences in DNA methylation at individual CpG sites across the methylome across 10 years in two Scandinavian samples of same-sex aging twins, estimating the genetic and environmental contributions to stability as well as to novel influences that emerge. Moreover, we examine whether surrounding ‘shores’ and ‘shelves’ are differentially heritable compared to islands, and, whether sites identified as associated with rate of aging in epigenome-wide association study (EWAS) or individual CpG clock sites are differentially heritable. In a combined sample of aging twins, assessed a decade apart in late-life, we test two competing hypotheses about the longitudinal stability and change in DNA methylation that stem from prior cross-sectional work (van Dongen et al., 2016): (1) the contribution of genetic influences changes with age, reflecting diminishing influence across time, and (2) nonshared factors accumulate in importance, signaling an increasing diversity of response to environmental exposures.

## Methods

### Sample

We considered DNA methylation across a 10-year span in 96 pairs of same-sex aging twins (40 monozygotic, MZ pairs; 56 dizygotic, DZ pairs). Across two samples, the average age at time 1 was 68.89 years (*S*D=8.58) and at time 2 was 78.59 years (*SD*=8.70). Specifically, the Swedish Adoption Twin Study of Aging (SATSA) included 53 pairs (22 MZ, and 31 DZ pairs; 53% female), selected with measurements about 10 years apart (range = 8.00 to 11.82 years) at ages 62.9 and 72.5 years at time 1 and time 2 respectively (SD=7.2). In 4 of 53 SATSA pairs, one twin partner had methylation data from one time point instead of both time points, but all data were included for these pairs. The Longitudinal Study of Aging Danish Twins (LSADT) included 43 pairs (18 MZ, and 25 DZ pairs; 72% female) at ages 76.2 and 86.1 years at time 1 and time 2 (SD=1.8).

### Materials

Methylation measurements from the Illumina HumanMethylation450 array (Illumina, San Diego, CA, USA) were preprocessed and normalized with adjustments for cell counts and batch effects. Processing of the SATSA sample probes has been described previously (Jylhävä et al., 2019; Y. Wang et al., 2018) and in brief included: (a) preprocessing with the R package *RnBeads* (Assenov et al., 2014) where filtering of samples and probes proceeded with a greedy-cut algorithm maximizing false positive rate versus sensitivity at a detection p-value of 0.05; (b) removal of sites that overlap with a known SNP site or reside on sex chromosomes; (c) normalization of data using *dasen* (Pidsley et al., 2013); (d) applying a Sammon mapping method (Sammon, 1969) to remove technical variance; (e) adjustment for cell counts (M. J. Jones, Islam, Edgar, & Kobor, 2017); (f) correction for batch effects using the ComBat approach in the *sva* package (Leek et al., 2012).

Processing of the LSADT data has been described previously (Svane et al., 2018) and in brief included: (a) preprocessing with the R-package MethylAid (van Iterson et al., 2014) where samples below quality requirements were excluded and probes with detection p-value>0.01, no signal, or bead count<3 were filtered out; (b) removal of probes with >5% missing values, removal of sites that reside on sex chromosomes or cross-reactive probes; (c) normalization and batch-correction using functional normalization(Fortin et al., 2014) with four principal components.

Although Beta-values are preferred for interpretation of methylation, Beta-value units were translated into M-values via a log2 ratio for improved distributional properties for the analysis of individual differences (Du et al., 2010). After performing the preprocessing steps, 390,894 probes remained for SATSA and 452,920 CpG sites remained for LSADT.

Altogether 368,391 sites were in common across the Swedish and Danish samples. After the described QC pre-processing in SATSA, 49 of 53 pairs had methylation data available for both members of each pair at both time points, while in 4 pairs one cotwin member had data at both time points while their twin partner had data at one timepoint but not both. After preprocessing, LSADT sample had methylation data represented for both cotwins at both timepoints among the 43 pairs.

### Filtering of sites post-analysis

We conducted additional filtering of probes where model-fitting results evidenced means or variances outside of expected values. Specifically, we filtered based on the typical range of M-values (c.f., Du et al., 2010), with expected mean values falling outside the range −6.25 to 6.25 for 1812 sites under either the ACE or ADE models at either timepoint. Likewise, we filtered based on expected standard deviations exceeding 1.5 under either the ACE or ADE models (Du et al., 2010) resulting in 9554 sites out of range under either the ACE or ADE models at either timepoint. The effective reduction is sites was from 368,391 to 358,836 after dropping 9555 unique sites from the analysis set.

### Analysis

Bivariate biometrical twin models of M-values were fitted to all available data across the pairs using full-information maximum likelihood (FIML), adjusting for centered age (centered at the average age across time = age − 74 years), sex (0=males, 1=females), and country (0=Sweden, 1=Denmark). Bivariate ACE and ADE Cholesky models evaluated the degree to which additive genetic (A), dominance or non-additive genetic (D), common environmental (C), and non-shared factors (E), encompassing non-shared environmental influences, measurement error, and stochastic factors, contributed to variation and covariation in M-values within and across time (see Figure 1). The resolution of the genetic and environmental effects are done by comparing the relative similarity of monozygotic (MZ) twins who share 100% of their genes in common, including all additive effects and dominance deviations, versus dizygotic (DZ) twins who share on average 50% of segregating genes in common leading to expectations of 50% for additive effects and 25% for dominance deviations. Both twin types are presumed to have the same contribution of common environmental effects that contribute to similarity. We fitted ADE and ACE models as dominance (D) and common environment (C) could not be simultaneously estimated (see Figure 1).

**Figure 1.**
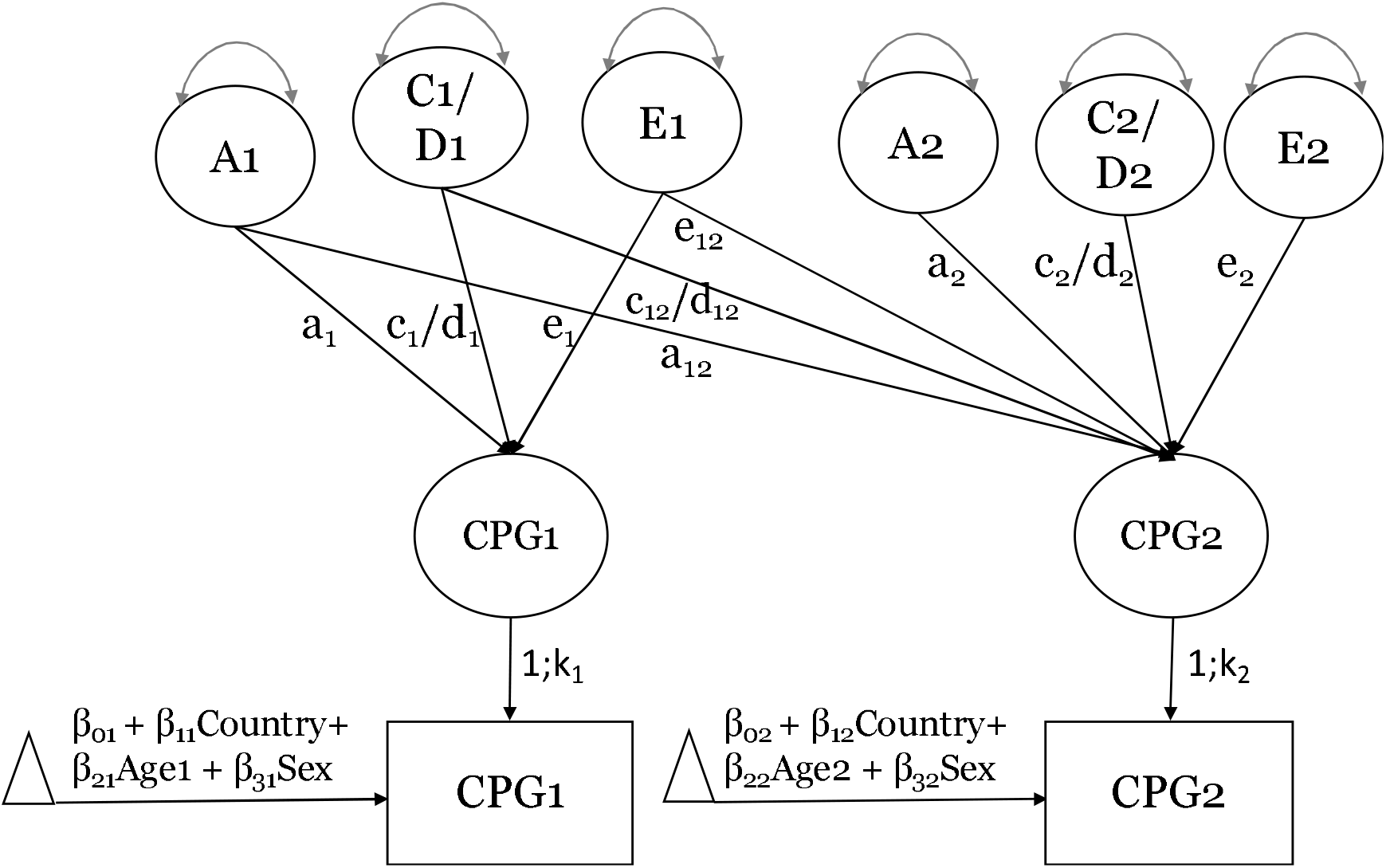
Bivariate Cholesky model. *Note.* ACE and ADE models were separately fitted to M-values at two waves 10 years apart.

Fit comparison between the ACE and ADE models was done via Akaike Information Criterion (AIC; Akaike, 1974). If the fit of the ADE model was as good or better than the ACE model it was retained as ‘best’ fitting, and otherwise the ACE model was retained as best. We evaluated submodels including AE, CE and E models. Differences in nested model deviance statistics [-2ln(L)] are distributed as chi-square (χ^2^) with the difference in the number of parameters between the full and constrained models as the degrees-of-freedom (df). LSADT samples tended to show lower variability in methylation at any given probe compared to SATSA, hence we allowed for scalar differences at each time-point (k_1_, k_2_) in standard deviations between the two samples (see Figure 1). Thus, the relative contributions of A, C or D, and E were equated across LSADT and SATSA, but the scalar allowed for the variance components to differ by a constant at each assessment. Scalar differences in standard deviations were on average k_1_ = .90 (SD=.93) and k_2_ = .88 (SD=.89).

Annotation of CpG sites with respect to UCSC CpG Island information (Gardiner-Garden & Frommer, 1987) was done by merging analysis results to the manifest file available for the Infinium HumanMethylation450 v1.2 BeadChip (Illumina, San Diego, CA, USA). Annotations included ‘Island’, ‘North’ ‘Shore’, ‘South’ ‘Shore’, ‘North’ ‗Shelf’, ‘South’ ‘Shelf’, and a blank annotation field was treated as ‘Open Seas’.

In comparing relative heritabilities across sites by location, as well as aging/clock CPGs sets to remaining CpGs, we fitted random effects regression models to age 69 and 79 biometrical estimates using *lme* (version 1.1-21; Bates, Mächler, Bolker, & Walker, 2015). We allowed for random effects between and within sites, reflecting consistency of effects by CpG sites across age and nonsystematic variation across time.

To compare time1-time2 correlations from the biometrical estimates we rescaled the a_12_, d_12_ or c_12_ and e_12_ paths into correlations (r_A_, r_D_ or r_C_, and r_E_) and performed Fisher Z-transformations before submitting each to a skew-normal regression analysis using the *sn* package (Azzalini, 2020). Regression analyses compared low stability sites to remaining CpGs, after which regression weights were inverse-transformed into correlation units for interpretation.

Enrichment analyses were conducted GREAT 4.0.4 (McLean et al., 2010). Selected sites were mapped to the Human GRCh37 build and default settings were used for association rules (i.e., basal+extension: 5000 bp upstream, 1000 bp downstream, 1000000 bp max extension, curated regulatory domains included). We present results of both biomial and hypergeometric tests where the False Discovery Rate (FDR) achieved *p* < .05 and where fold enrichment (FE) tests exceeded 2.0. We followed up the enrichment analyses using the mQTL Database (Gaunt et al., 2016) to annotate associations with methylation quantitative trait loci, noting the number of *cis* or *trans* variants.

## Results

We first evaluated the extent to which heritable and environmental influences contributed to each CpG site. Bivariate biometrical twin model results, comparing MZ twin similarity to DZ twin similarity within and across time, suggests under an ADE model that broad-sense heritable contributions (A+D, N=358,836) were on average small at age 69 years (*M* = 0.238*100 = 23.8%, time 1) and decreased across 10 years (*M* = 0.180*100 = 18.0%, time 2) (see Table 1, Variance Components). The decrease in broad heritability across time is significant within site, *M*_t2-t1_ = −.058 (*t* = −232.0, df=358835, CI_95_ = −.058, −.057). The decrease in heritability is due to an absolute increase in non-shared factors (E) compared to genetic influences (A, D) (see Table 1, Absolute Variances). Patterns of decline were observed for heritabilities (A) under the ACE model (. 150 and .109, respectively), and under best-fitting ADE or ACE models (see Table 1, Variance Components). Common environmental influences were generally stable in overall ACE results at over 5% (.057, .054) and in best-fitting ACE results at 10% (.106, .098) (see Table 1, Variance Components).

**Table 1.** Variance components and absolute variances at time 1 (69 years) and time 2 (79 years).

| **Variance Components** | **N sites** | **A1** |  | **D1/C1** |  | **E1** |  | **A2** |  | **D2/C2** |  | **E2** |  |
| --- | --- | --- | --- | --- | --- | --- | --- | --- | --- | --- | --- | --- | --- |
|  |  | *M* | *SD* | *M* | *SD* | *M* | *SD* | *M* | *SD* | *M* | *SD* | *M* | *SD* |
| ADE | 358,836 | 0.111 | 0.142 | 0.127 | 0.160 | 0.762 | 0.175 | 0.091 | 0.125 | 0.089 | 0.130 | 0.820 | 0.158 |
| ADE best | 187,535 | 0.057 | 0.106 | 0.217 | 0.168 | 0.725 | 0.182 | 0.048 | 0.092 | 0.152 | 0.148 | 0.800 | 0.168 |
| ACE | 358,836 | 0.150 | 0.166 | 0.057 | 0.083 | 0.793 | 0.163 | 0.109 | 0.142 | 0.054 | 0.080 | 0.837 | 0.147 |
| ACE best | 171,301 | 0.076 | 0.122 | 0.106 | 0.093 | 0.817 | 0.150 | 0.055 | 0.103 | 0.098 | 0.091 | 0.846 | 0.137 |
| **Absolute Variances** | **N sites** | **A1** |  | **D1/C1** |  | **E1** |  | **A2** |  | **D2/C2** |  | **E2** |  |
|  |  | *M* | *SD* | *M* | *SD* | *M* | *SD* | *M* | *SD* | *M* | *SD* | *M* | *SD* |
| ADE | 358,836 | 0.029 | 0.079 | 0.038 | 0.088 | 0.162 | 0.144 | 0.028 | 0.079 | 0.032 | 0.086 | 0.214 | 0.195 |
| ADE best | 187,535 | 0.019 | 0.063 | 0.067 | 0.112 | 0.174 | 0.151 | 0.018 | 0.063 | 0.057 | 0.111 | 0.235 | 0.206 |
| ACE | 358,836 | 0.047 | 0.109 | 0.012 | 0.034 | 0.170 | 0.152 | 0.042 | 0.109 | 0.014 | 0.038 | 0.220 | 0.200 |
| ACE best | 171,301 | 0.021 | 0.074 | 0.023 | 0.045 | 0.152 | 0.137 | 0.019 | 0.072 | 0.025 | 0.050 | 0.193 | 0.180 |

*Note.* A = Additive genetic, D = Nonadditive genetic (Dominance), C = Common environment, E = Non-shared factors.

Across time, heritabilities showed divergence by location [ADE best (A+D): χ^2^ (5) = 618.3, *p* = 2.25E-131; ACE best (A): χ^2^ (5) = 339.5, *p* = 3.19E-71] (see Table S1, Figure 2). In ADE best results, islands and shelves showed lower broad (A+D) heritabilities than open seas by −.01 or −1% (*p* ≤ 1.55E-07) whereas shores were higher by .01 or 1% than open seas (*p* ≤ 3.79E-15). In ACE best results, comparably lower heritabilities (A) were observed for islands versus open seas (*p* = 3.54E-58).

**Figure 2.**
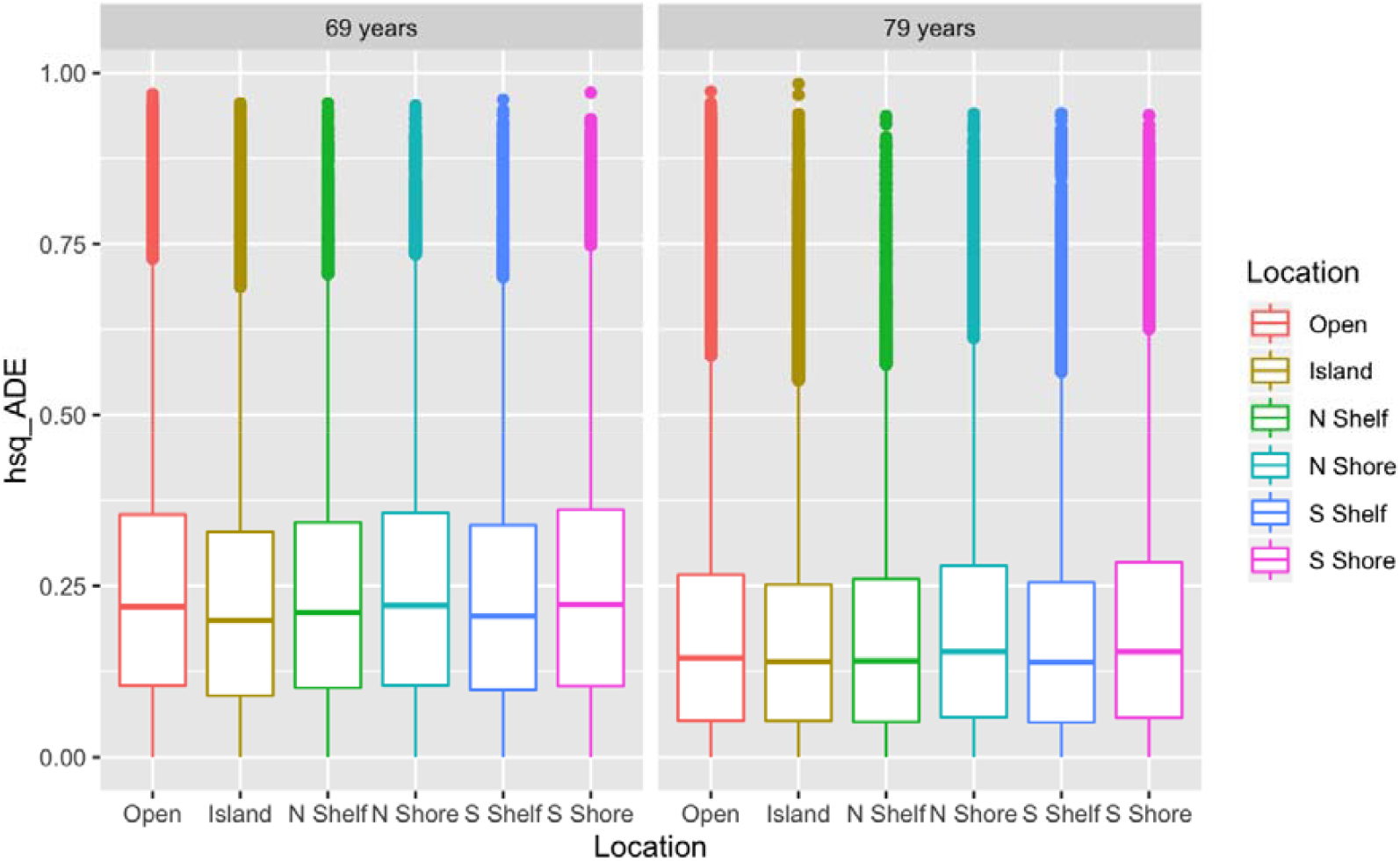
Broad-sense heritability by Location across 10 years (ADE results, 358,836 CpGs) *Note.* Site differences shown below are significant across time: χ^2^(5)=995.48, p=5.72E-213

Next, we evaluated the number of CpG sites that achieved significant heritable or familial-environmental effects. At epigenome-wide significance (*p*<1E-07), 5037 CpG sites (1.4%) showed broad genetic (A, D) or familial-environmental effects (A, C) within or across time (*df*= 6), and 35,762 sites (10.0%) met *p*<1E-02. Among the 358,836 sites, 52% of sites showed the better-fitting model was ADE (*N*=187,535) while 48% showed ACE as better-fitting (*N*=171,301) (see Table 1, Figure 3). A total of 58,676 sites (16.4%) achieved nominal significance comparing the ADE or ACE versus an E model (*p*<.05, 6 df; *N*=32685 ADE best, *N*=25991 ACE best), and 91,380 sites (25.5%) achieved nominal significance of an AE model over an E model (*p*< .05, df=3). Given that power is low for C even in large samples, as well as to distinguish D from A, we present full model estimates (Visscher, Gordon, & Neale, 2008).

**Figure 3.**
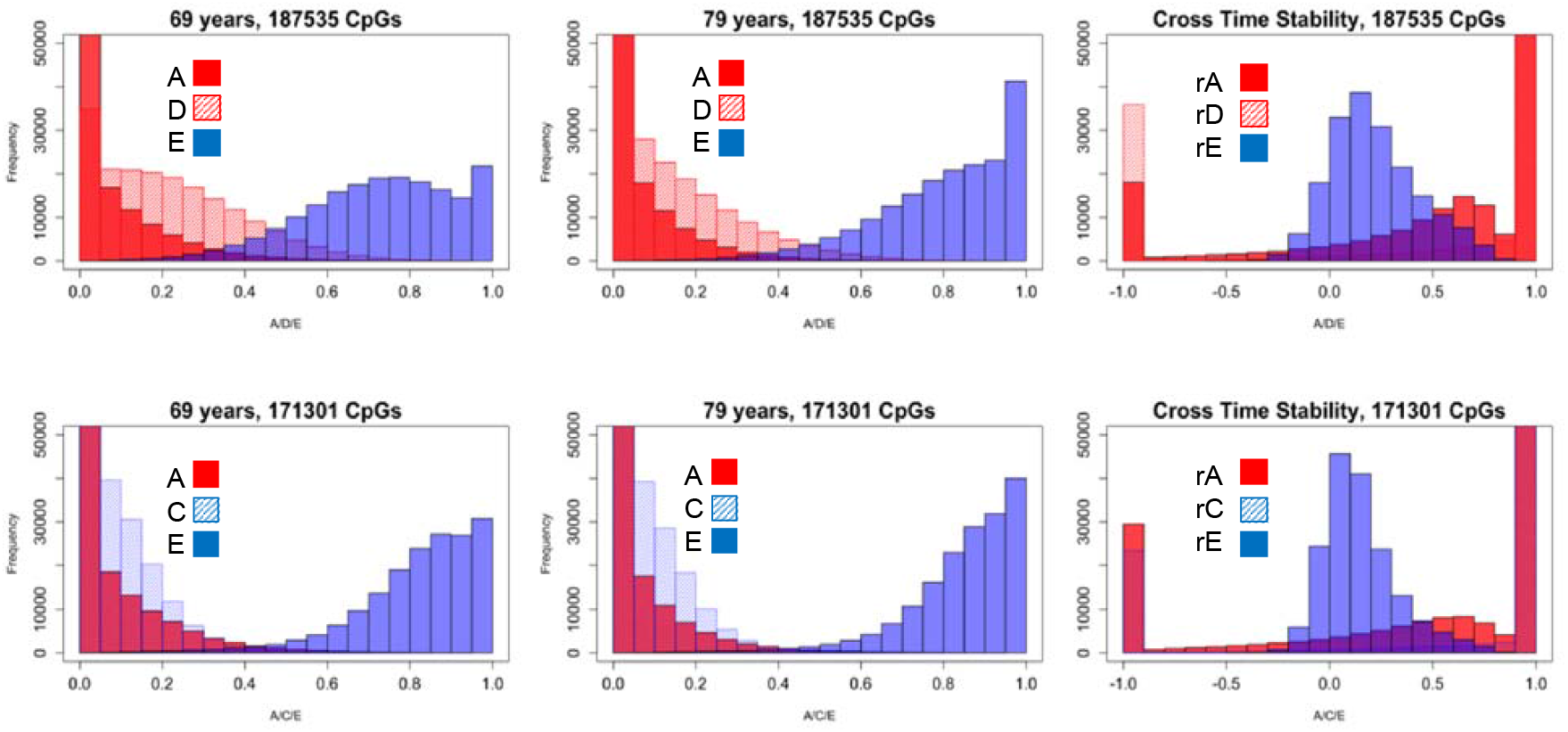
Best-fitting models: ADE (52%) or ACE (48%)

**Figure 4.**
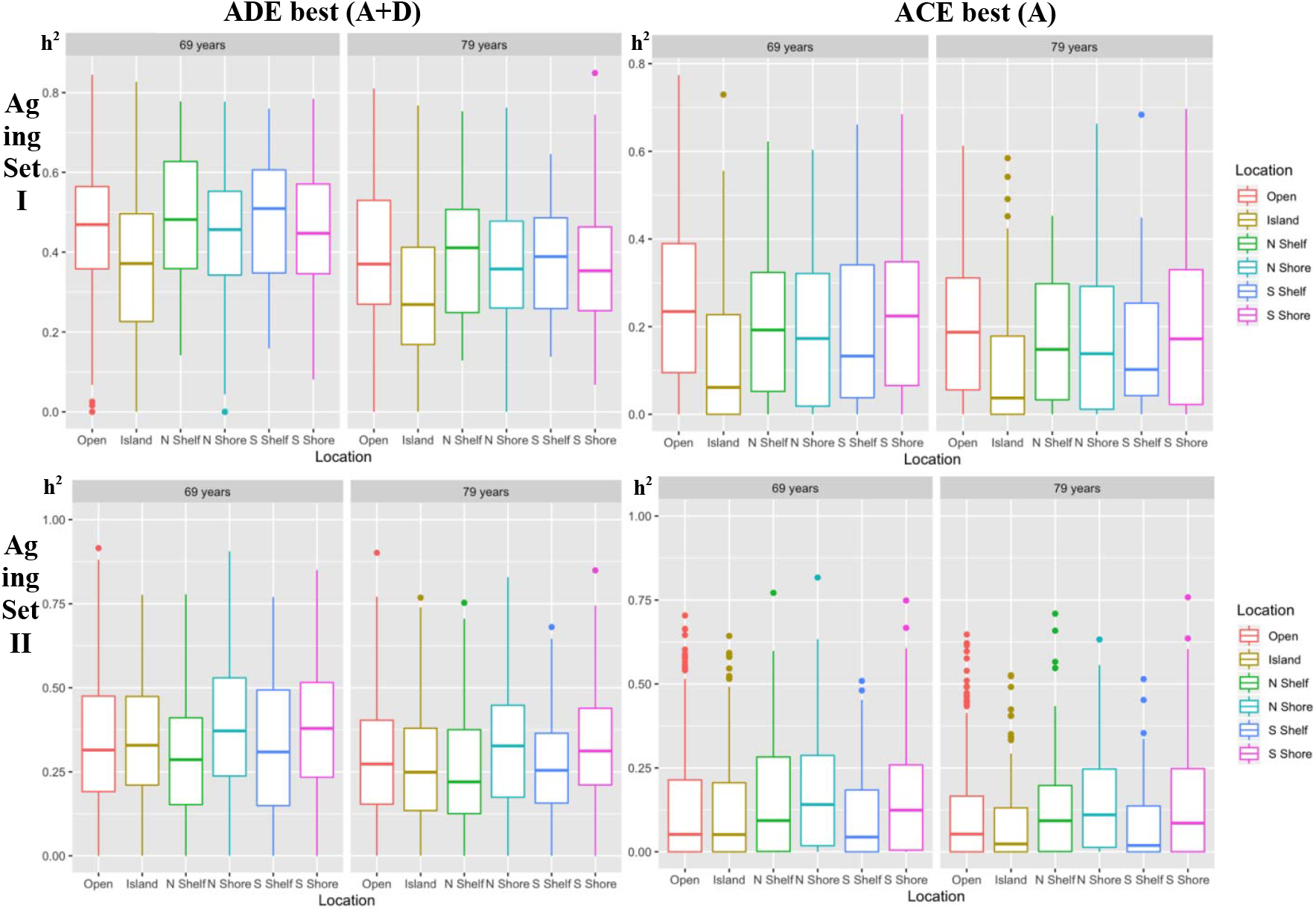
Age-related CpG Sets: Broad heritability by CpG Location

In terms of contributions to stability and change in methylation due to genetic or environmental influences, across 358,836 sites, 58.5% showed cross-time associations at p<.05 (df = 3, where a_12_=[d_12_ or c_12_]=e_12_=0) indicative of stability over time due either to genetic and/or environmental mechanisms. As shown in Figure 3, the cross-time stability was largely due to genetic effects in both the ADE best and ACE best models which was most often perfect in correlation.

As cross-sectional twin studies have reported that heritability may be higher for variable methylated sites (e.g., Hannon et al., 2018), we report the correlation between the estimated standard deviations of M-values and the extent to which heritable effects were observed at time 1 and 2, respectively: (1) *r_SD,A+D_* = .33 and .27 (187535 sites) for ADE best, and (2) *r_SD,A_* = .25 and .20 (171301 sites) for ACE best. Sites in which nonshared factors, E, explained all of the variability of M-values (>99%) at both time points included 8,268 total sites (5,520 ADE best, 2,748 ACE best). In all these cases, we observed that either the MZ twin correlations of M-values were negative (< 0), or DZ correlations were sufficiently negative (< −.05), or the difference between MZ and DZ correlations at each time point were sufficiently negative (< −.1).

### Age-related sites

We evaluated the best-fitting ADE and ACE results of two published CpG sets that were identified in EWAS as related to Age that overlap with the samples used in the presented analysis: (I) 1217 sites from Wang et al. (Y. Wang et al., 2018); (II) 1934 sites from Tan et al. (Tan et al., 2016). Multilevel regression models compared heritabilities by location from the ADE best or ACE best model, fitted to both age 69 and 79 estimates in set I [ADE best (A+D): χ^2^ (5) = 43.7, *p* = 2.66E-08; ACE best (A): χ^2^ (5) =27.9, *p* = 3.81E-05], with Islands under ADE or ACE models showing lower heritabilities by .09-.10 or up to a 10% difference than open seas (both *p* ≤ 5.13E-07; see Table S1). In set II, age 69 and 79 heritability estimates also showed divergence by location [ADE best (A+D): χ^2^ (5) 16.8, *p* = 4.90E-03; ACE best (A): χ^2^ (5) = 19.4, *p* = 1.62E-03], with Shores showing higher heritabilities by about .04 or 4% than open seas under ADE or ACE models (all *p* ≤ 2.56E-02; see Table S1).

Multilevel regression models were fitted to age 69 and 79 biometrical estimates to compare the Aging sets’ CpG sites to the remaining background CpGs. Stronger heritable influences were apparent for Aging Set I (1217 sites) compared to remaining CpGs with .16 higher broad heritability (.39 vs .24, ADE best; *p* = 5.43E-162) and .12 higher narrow heritability (.18 vs .07, ACE best; *p* = 3.87E-146) and .05 higher common environmentality (.15 vs .10; *p* = 1.88E-40) (see Table S2, Variance Components). Patterns in the absolute variances suggested the greater heritability was due primarily to lower nonshared factors (*p* ≤ 1.35E-07) and for ACE models coupled with higher additive genetic and common environmental influences (*p* ≤ 2.03E-04; see Table S2, Absolute Variances). Significantly higher heritabilities and common environmentality were also observed for Aging Set II (1934 sites) where the increased heritable and common environmental influences (all *p* ≤ 1.55E-31) were driven mainly by amplified genetic and common environmental influences *(p* ≤ 1.01E-11) and otherwise comparable non shared factors between the Aging II set and remaining background CpGs (see Table S2). Thus, the age-related sites showed a significantly higher proportion of variance attributed to heritable and shared environmental influences due to lower nonshared factors in Aging set I and due to higher genetic and common environmental influences in Aging set II.

### Methylation clock sites

Available CpG sites from four epigenetic-clocks were evaluated in similar fashion using multilevel regression models fitted to age 69 and 79 biometrical estimates: (a) 59 of 71 sites Hannum clock (Hannum et al., 2013), (b) 312 of 353 sites Horvath clock (Horvath, 2013), (c) 443 of 513 sites Levine clock (Levine et al., 2018), and 455 of 514 sites from the Zhang clock (Q. Zhang et al., 2019). Significantly higher heritabilities (A+D, A) and common environmentality (C) were observed for the 1190 unique Clocks sites compared to all remaining CpGs (.02 to .05 higher, *p* ≤ 8.68E-10, see Table S2 Variance Components). Comparisons of absolute variances suggested amplified genetic and common environmental influences (*p* ≤ 3.94E-05) as well as non-shared factors (*p* ≤ 3.90E-08) between the Clock sites and remaining background CpGs (see Table S2, Absolute Variances). Thus, the clock sites showed greater overall variability across sources of variance suggesting greater individual differences in these sites, with a significantly higher portion of variance attributed to heritable and shared environmental influences.

Among the 1190 unique CpG sites compared to one another, the Zhang clock sites tended to show stronger broad (A+D) genetic (.07 to .08 higher, *p* ≤ 4.00E-07), and shared environmental (C) contributions (.06 higher, *p* ≤ 2.19E-08) than Horvath or Levine clock sites, while Hannum sites were comparable to Zhang sites (within −.018 to .018, *p* > 4.71E-01) (see Table S3). The ratio of Intercept variance to total variance (*ρ*) in heritability estimates was .559 for ADE best models and .680 for ACE best models suggesting 56% and 68% of the variation in heritability, respectively, was CpG site specific across time and less than half of the variation was unique to CpG site and time, consistent with analyses by location (see Table S1). Likewise, absolute variances showed strong between-site variations (66%-88%, see Table S3).

### Low Stability Sites

We identified 2020 CpGs with low stability but meaningful genetic or common environmental contributions at one or both timepoints, i.e., *p* > .01 (df = 3, where a_12_=[d_12_ or c_12_]=e_12_=0) and where e_1_ or e_2_ accounted for less than 50% of the total variation (1638 ADE best, 382 ACE best). Based on skew-normal analyses, low stability CpGs had lower correlations among non-shared factors across time than background CpGs (ADE best: *r*_E,background_ = .24 vs *r*_E,low_ = .10, *p* = 2.80E-154; ACE best: *r*_E,background_ = .17 vs *r*_E,low_ = .07, *p* ≤ 3.09E-27). The correlations of genetic (r_A_, r_D_) and common environmental influences (r_C_) across time were comparable (within .02 units) between background and low stability CpGs, albeit significant *(p* ≤ 1.02E-03), and otherwise very strong based on skew-normal analyses (*r*_background_ = 97 − .99 vs *r*_low_ = .95 − .99). Variability in these low stability CpGs increased across time with a ratio of SD_2_/SD_1_ of 1.08 to 1.09 (SD_ratio_ = 13) for ACE and ADE best models, respectively. Moreover, heritabilities decreased across time while non-shared components tended to increase (see Figure S1). Compared to background CpGs, low stability CpGs tended to show higher A+D or A and C components (all *p* ≤ 2.03E-14) but generally lower overall absolute variances for A+D and E variances *(p* ≤ 15.51E-09) in ADE models (see Table S2). Higher absolute variance for A but lower variance for E was observed in ACE models (*p* ≤ 4.94E-03) (see Table S2). Altogether, results suggest lower overall phenotypic variance in methylation among the low stability versus background CpGs across time (c.f., Table S2). However, within the set of lower stability CpGs, variance in methylation increased at time 2 mainly due to novel non-shared factors (c.f., Figure S1).

### Enrichment analysis: High heritabiliy/familialiy

The set of 5037 CpGs achieving epigenome significance (*p*<1E-07) when evaluating tests of heritability (AD vs E; N = 2049) or familiality (AC vs E; N=2988) across time were submitted to GREAT 4.0.4 to identify functions of cis-regulatory regions (McLean et al., 2010). Specifically, we report the binomial and hypergeometric tests over genomic regions covered by the 5037 CpGs, reporting those that achieved region-based fold enrichment (FE) > 2 and both binomial and hypergeometric FDR Q-Values < .05 (see Table 2; for full ontology results see Table S4). The sites that showed the greatest heritabilities showed enrichment in immune and inflammation pathways as well as neurotransmitter activity pathways. For example, the MHC protein complex pathway in the GO Cellular ontology list includes HLA region genes that code for HLA class II histocompatibility antigens in humans (c.f., GO:0042611, Table S4). Moreover, the interferon-gamma-mediated signaling pathway in the GO Biological ontology list include numerous genes associated with altered cytokine signaling and genes in the HLA region (c.f., GO:0060333, Table S4).

**Table 2.**
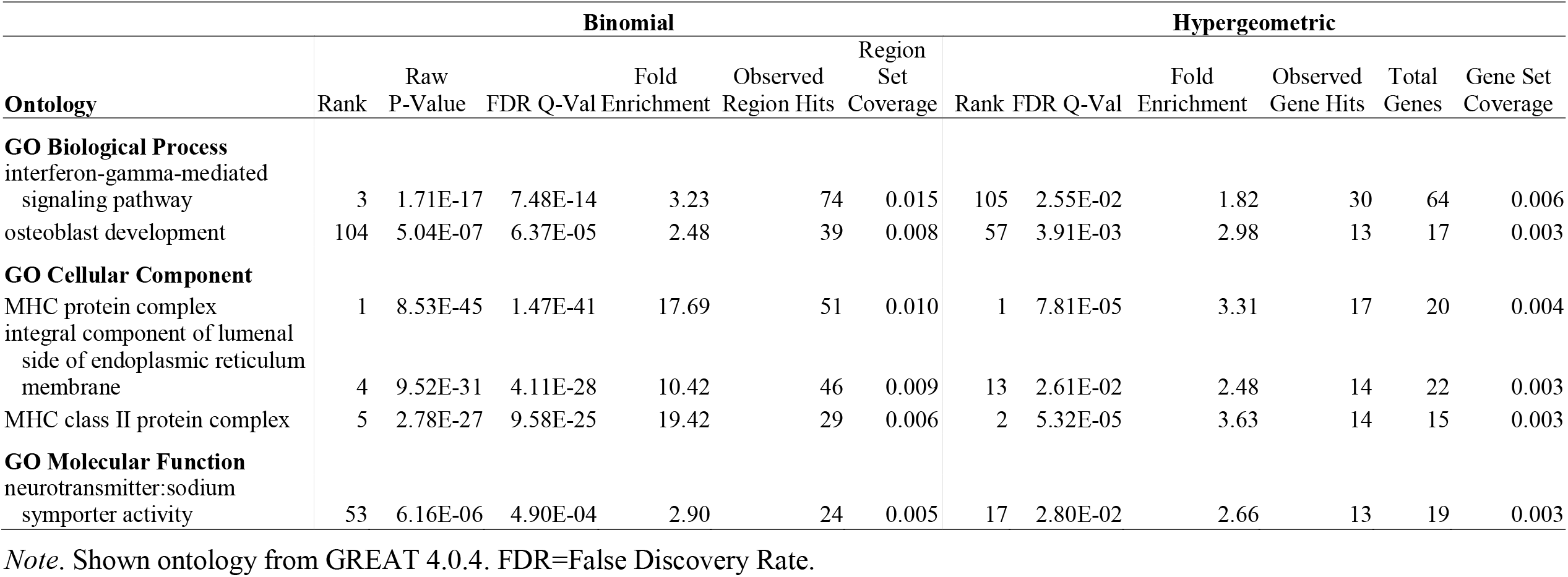
GREAT 4.0.4 annotations using binomial and hypergeometric tests over genomic regions covered by the 5037 CpGs showing significant heritability/familiality p<1E-07.

| Ontology | Binomial |  |  |  |  |  | Hypergeometric |  |  |  |  |  |
| --- | --- | --- | --- | --- | --- | --- | --- | --- | --- | --- | --- | --- |
|  | Rank | Raw P-Value | FDR Q-Val | Fold Enrichment | Observed Region Hits | Region Set Coverage | Rank | FDR Q-Val | Fold Enrichment | Observed Gene Hits | Total Genes | Gene Set Coverage |
| **GO Biological Process** |  |  |  |  |  |  |  |  |  |  |  |  |
| interferon-gamma-mediated signaling pathway | 3 | 1.71E-17 | 7.48E-14 | 3.23 | 74 | 0.015 | 105 | 2.55E-02 | 1.82 | 30 | 64 | 0.006 |
| osteoblast development | 104 | 5.04E-07 | 6.37E-05 | 2.48 | 39 | 0.008 | 57 | 3.91E-03 | 2.98 | 13 | 17 | 0.003 |
| **GO Cellular Component** |  |  |  |  |  |  |  |  |  |  |  |  |
| MHC protein complex | 1 | 8.53E-45 | 1.47E-41 | 17.69 | 51 | 0.010 | 1 | 7.81E-05 | 3.31 | 17 | 20 | 0.004 |
| integral component of luminal side of endoplasmic reticulum membrane | 4 | 9.52E-31 | 4.11E-28 | 10.42 | 46 | 0.009 | 13 | 2.61E-02 | 2.48 | 14 | 22 | 0.003 |
| MHC class II protein complex | 5 | 2.78E-27 | 9.58E-25 | 19.42 | 29 | 0.006 | 2 | 5.32E-05 | 3.63 | 14 | 15 | 0.003 |
| **GO Molecular Function** |  |  |  |  |  |  |  |  |  |  |  |  |
| neurotransmitter:sodium symporter activity | 53 | 6.16E-06 | 4.90E-04 | 2.90 | 24 | 0.005 | 17 | 2.80E-02 | 2.66 | 13 | 19 | 0.003 |

*Note.* Shown ontology from GREAT 4.0.4. FDR=False Discovery Rate.

The set of 5037 CpGs were then submitted to the mQTL Database (Gaunt et al., 2016). The search resulted in 1435 unique CpG matches to 155,177 SNP variants from the Middle Age timepoint (see Table S6). Of the 1435 CpG matches, 1256 were associated with cis-mQTLs and 304 were associated with trans-mQTLs suggesting an abundance of associations with cis-mQTLs. The maximum number of mQTLs associated with any given CpG was for cg03202060 with 5230 cis-mQTLs variants plus 575 trans-mQTLs. The cis-mQTLs for cg03202060 reside in the HLA region on chromosome 6 (e.g., https://www.genecards.org/cgi-bin/carddisp.pl?gene=HLA-DQB1&keywords=HLA-DQB1), and the trans-mQTLs traverse genes such as *DDAH2* related to metabolism of nitric oxide (https://www.genecards.org/cgi-bin/carddisp.pl?gene=DDAH2) and *BAG6* (https://www.genecards.org/cgi-bin/carddisp.pl?gene=BAG6&keywords=BAG6) residing within the major histocompatibility class III region (MHCIII) and involved in the control of apoptosis. A scatterplot of cg03202060 M-values of twin 1 by twin 2 across time is shown in Figure S2a,b showing greater similarity for MZ than DZ pairs.

As polycomb repression may relate to age-related changes in DNA methylation, we filtered our set of 5037 CpGs to reflect genes annotated on the 27k array and evaluated whether our set mapped to 1861 PolyComb Group Target genes (PCGTs) identified using the Illumina 27k chip probes (Zhuang et al., 2012). We observed 493 CpGs within a set of 293 PCGTs overlapped, or a 15.7% overlap of PCGTs (see Table S6). A hypergeometric test of the 293 overlapping PCGTs was significant at *p* = 1.004E-11 suggesting overrepresentation, when considering the number of unique PCGTs in Zhuang et al. (2012), and the number of genes represented in the Illumina 27k chip.

### Enrichment analysis: Low stability sites

The 2020 low stability CpGs were submitted to GREAT 4.0.4, showing enrichment for stress-related DNA and RNA transcription pathways (see Tables S7-S8). Hence, these sites may lie in genes/gene pathways that are sensitive to exogenous exposures to stress leading to increasing divergence in methylation profiles across time. The GO Biological RNA and DNA pathways noted relate to heat shock and response to hypoxia in a number of plant and animal species, including humans (c.f., annotations GO:0043620, GO:0061418; Table S8).

The low stability CpGs were submitted in kind to the mQTL Database (Gaunt et al., 2016) producing 397 unique CpG matches to 7103 mQTLs at the Midlife timepoint. Of the 397 CpG matches, 58 annotations were to cis-mQTLs and 347 were to trans-mQTLs (see Table S9), suggesting an abundance of associations with trans-mQTLs. The maximum number of mQTLs linked with any given CpG was for cg07677296 matched with 576 cis-mQTLs. The cis-mQTLs variants associated with cg07677296 traverse *FAHD1* and *NUBP2* on chromosome 16 and have been implicated in aging pathways related to insulin-like growth factor (Teumer et al., 2016). A scatterplot of cg07677296 M-values of twin 1 by twin 2 across time shows comparable similarity for MZ and DZ pairs (see Figure S2c,d).

## Discussion

Overall, results suggest genetic contributions to DNA methylation tended to be small, vary by location, and decrease across a decade; however, genetic influence mainly contributed to the stability of methylation. Unique person-specific influences not shared by co-twins were emergent across 10 years suggesting that non-shared factors become more salient to DNA methylation in late life. The extent of variation in methylation at any given CpG site was positively correlated with observing stronger heritable effects. Moreover, 58% of sites showed stability across time due to strongly correlated genetic influences and modestly correlated nonshared factors, suggesting continuity of influences across 10 years for more than half the CpG sites. The sites that showed the greatest heritabilities showed enrichment in immune and inflammation pathways and neurotransmitter transporter activity pathways. Low stability sites meanwhile showed increased expression variability across time due to novel nonshared factors, with enrichment in stress-related pathways, suggesting that these sites are responsive to “new” environmental cues even in old age.

Prior studies report average heritabilities of 16.5 – 19.0% across adulthood (17-79 years) (Hannon et al., 2018; van Dongen et al., 2016) and common environmental influences of 3.0 − 12.6%, that are stronger in young adulthood (Hannon et al., 2018). Our results of weakening heritable influences across age is consistent with the Dutch cross-sectional study reporting average heritabilities of 21% and 18% at ages 25 and 50 assuming an AE model (van Dongen et al., 2016), whereas our estimates of broad heritability under an ADE model are 24% and 18% many decades later at age 69 and 79 years, respectively. Where non-additive genetic effects fit best, the average broad heritability was 24% across age. For sites where including common environment fit best (ACE), lower average heritabilities were observed at 7% whereas common environment contributed 10% to variation in methylation across age; common environment is higher in 18-year-old UK adults at 12.6% (Hannon et al., 2018). We directly compared our heritabilities with those available from Van Dongen et al. (2016) where twins were on average 37.2 years (17-79 years). For 337,322 matching sites, our A+D estimates at time 1 (69 years) were strongly correlated with their AE results (*r* = .568, df = 337,320, CI_95_ = 0.566, 0.570) and with their total heritability estimates where age interactions were estimated (r = .556, df= 334,657, CI_95_ = 0.554, 0.559).

CpG sites related to age show a greater impact of heritable influences consistent with genetic regulation of the rate of biological aging. Sites associated with age and longevity generally show higher heritabilities than the total background sites and varied in magnitude of heritabilities by location, where ‘islands’, which often reside in promotor regions (Vinson & Chatterjee, 2012), typically showed lower heritability than those sites residing in surrounding ‘shores’ and ‘shelves’, which have been shown to be differentially methylated compared to islands (M. J. Jones et al., 2015).

Moreover, the set of methylation clocks sites are likewise more heritable than background CpG sites, with Zhang sites more heritable than Horvath and Levine sites, and Hannum sites comparable to the Zhang sites. We have recently reported heritability estimates of methylation clock ages of 52% for the Horvath clock and 36% for the Levine clock (Jylhava et al., 2019), where, consistent with our current site-specific effects, stability across time was mediated primarily by genetic factors, whereas the person-specific environmental factors contributed to differences across time. The 353 Horvath clock sites were selected as best predictors of chronological age using multiple tissues (Horvath, 2013) similar to the 513 Levine clock sites that were selected based on prediction of chronological age and nine biomarkers of phenotypic aging with models trained on multiple tissues (Levine et al., 2018). The 71 Hannum clock sites best predicted age (adjusted for sex, BMI) based on methylation observed in whole blood while the 514 sites from the Zhang prediction model relied on methylation observed in blood and saliva samples (Q. Zhang et al., 2019). The current findings of moderately higher heritabilities in the Zhang and Hannum sites versus the other clock sites may be in part due to our use of blood tissue.

Enrichment analyses of the 1.4% of sites meeting *p*<1E-07 suggest immune and inflammation pathways and neurotransmitter transporter activity pathways may feature in sites with strong heritable or familial-environmental components. Moreover, the analysis of mQTL associations suggest that a number these high heritability CpGs are associated largely with cis-mQTLs, including those in the HLA region. Previous studies have identified methylation changes associated with altered immune functioning, including age-related hypermethylation and reduced expression in CD8+ cells for genes involved in T cell mediated immune response and differentiation (Tserel et al., 2015). Indeed, five CpGs in our set identified as associated with cis-mQTLs at midlife, lie within the *BCL11* gene (cg26396443) or *RUNX3* gene (cg05162523, cg13566436, cg20674490, cg22509179) involved in T cell differentiation (Tserel et al., 2015). A related study of German and Danish individuals (including an overlapping sample of twins herein) evaluating RNA-sequencing expression patterns and longevity identified expression patterns in biological processes contributing to immune system and response pathways (Häsler et al., 2017), and observed high heritabilities (30-99%) among 20% of cis-eQTLS. Immunosenescence describes an age-associated decline in elderly individuals’ immune functioning, such as mounting less effective responses to vaccines, and lowered resistance to illnesses, with concomitant up-regulation of pro-inflammatory cytokines, among several other cellular and physiological changes in the immune system (Accardi & Caruso, 2018). It has been proposed that heritable factors may be partly associated with differential immune responses (Derhovanessian et al., 2010; Poland et al., 2014) and may predict influenza-related susceptibility and mortality (Poland et al., 2014), for example, and, broadly, successful aging and longevity (Derhovanessian et al., 2010). Hence, differential adaptions to aging processes including immunosenescence reflect gene-environment dynamics with some individuals showing better adaptions than others due to genetic influences.

High heritability CpGs were also enriched for PCGTs – a group of genes that are epigenetically regulated by polycomb-group proteins and involved in developmental processes and cell-fate decisions (Lanzuolo & Orlando, 2012). Enrichment of hypermethylated of PCGT has also been implicated in cancer and aging and show consistent patterns across different cell types (Teschendorff et al., 2010). Our findings would thus support the role of heritable/familial-environmental factors in the epigenetic regulation of these fundamental cellular processes.

Enrichment analyses of low stability CpG sites suggest that stress-related DNA and RNA transcription pathways may be relevant for these environmentally responsive sites which showed increased novel environmental contributions to methylation. It is notable that unlike the high heritability set, the low stability set showed more associations with trans-mQTLs. That said, cg07677296 matched with 576 cis-mQTLs, with variants spanning *FAHD1* and *NUBP2,* both implicated in metabolic and aging pathways related to insulin-like growth factor (IGF) (Teumer et al., 2016). Specifically, *FAHD1* was identified as a cis-eQTL associated with a variant in *NUBP2* (rs1065656) that may contribute to circulating IGF-I and IGFBP-3 concentrations (Teumer et al., 2016). Moreover, IGF-I is implicated in oxidative stress pathways (Gubbi et al., 2018).

The current study establishes the extent to which the genetic and environmental influences contribute to site-specific methylation across a 10-year span in a longitudinal sample of Swedish and Danish twins. While stability of methylation was largely due to genetic influences, person-specific environmental influences were emergent across time and explained change. By and large, the dynamics of methylation may be influenced by experiences and exposures, suggesting possible mediation of gene expression; however, the most heritable sites may participate in immune and inflammation pathways and neurotransmitter transporter activity pathways which suggest that adaptions to aging and senescence may be differentially impacted by genetic background.

## Funding

SATSA has been supported by the National Institute on Aging (AG04563, AG10175), the MacArthur Foundation Research Network on Successful Aging, the Swedish Council for Working Life and Social Research (FAS) (97:0147:1B, 2009-0795), and the Swedish Research Council (825-2007-7460, 825-2009-6141). DNA extraction was partly supported by AG028555, AG17561. Methylation work was supported by FORTE 2013-2292, the Swedish Research Council (521-2013-8689, 2015-03255) and a Distinguished Professor Award from the KI to NLP.

LSADT has been supported by grants from the VELUX FOUNDATION, the U.S. National Institute on Aging (P01-AG08761), the European Union’s Seventh Framework Programme (FP7/2007-2011) under grant agreement no. 259679, and The Danish National Program for Research Infrastructure 2007 [09-063256].

Collaborative work was supported in part by the National Institute on Aging (AG037985). The manuscript content is solely the responsibility of the authors and does not necessarily represent the official views of the National Institutes of Health.

## Conflict of Interest

none declared

## Author contributions

CAR drafted the manuscript. CAR and EM analyzed data. QT and JH contributed to scripting and QT, SH, and JJ advised on enrichment analyses. NLP and SH contributed to the coordination of the study and acquisition of the SATSA methylation data. JJ contributed to preparation of SATSA data and interpretation of results. LC, QT and JH coordinated the LSADT data acquisition. All authors participated in interpretation of the data, have read and commented on the manuscript and approved the final version.

## Notes

### Competing Interest Statement

The authors have declared no competing interest.

### Summary of Updates

Minor revisions made to text. Table 1 and Table S3 updated.

