## Supplemental Tables S1-S3_FigS1-S2 for "A decade of epigenetic change in aging twins: genetic and environmental contributions to longitudinal DNA methylation"

**Table S1. Multi-level regression results of variance components by location across age 69 and 79 years.**

| **ADE best (A+D)** | **Total sites (187,535)** | | | | **Aging I (717)** | | | | **Aging II (1,139)** | | | |
| --- | --- | --- | --- | --- | --- | --- | --- | --- | --- | --- | --- | --- |
| **Fixed effects** | **B** | **se** | **t** | **p** | **B** | **se** | **t** | **p** | **B** | **se** | **t** | **p** |
| Intercept  (Open Seas) | 0.239 | 0.001 | 393.80 | 0.00E+00 | 0.422 | 0.011 | 38.52 | 0.00E+00 | 0.313 | 0.008 | 37.25 | 9.29E-304 |
| Island | -0.012 | 0.001 | -13.30 | 2.24E-40 | -0.090 | 0.016 | -5.78 | 7.56E-09 | -0.007 | 0.013 | -0.55 | 5.84E-01 |
| N. Shelf | -0.009 | 0.002 | -5.25 | 1.55E-07 | 0.017 | 0.026 | 0.65 | 5.14E-01 | -0.026 | 0.023 | -1.14 | 2.55E-01 |
| N. Shore | 0.009 | 0.001 | 7.86 | 3.79E-15 | -0.018 | 0.017 | -1.05 | 2.95E-01 | 0.040 | 0.015 | 2.60 | 9.32E-03 |
| S. Shelf | -0.011 | 0.002 | -6.11 | 9.79E-10 | 0.004 | 0.032 | 0.12 | 9.04E-01 | -0.015 | 0.024 | -0.61 | 5.41E-01 |
| S. Shore | 0.013 | 0.001 | 10.71 | 8.69E-27 | -0.011 | 0.017 | -0.64 | 5.21E-01 | 0.035 | 0.016 | 2.23 | 2.56E-02 |
| **Random Effects** | **Intercept** | **Resid** | **ρ** | **--** | **Intercept** | **Residual** | **ρ** | **--** | **Intercept** | **Residual** | **ρ** | **--** |
| Variance | 0.016 | 0.016 | 0.505 | -- | 0.017 | 0.012 | 0.577 | -- | 0.023 | 0.013 | 0.635 | -- |
| **Model compare** | **Equal** | **Full** | **Δ χ2 (5)** | **p** | **Equal** | **Full** | **Δ χ2 (5)** | **p** | **Equal** | **Full** | **Δ χ2 (5)** | **p** |
| Deviance | -281864.9 | -282483.2 | 618.3 | 2.25E-131 | -1243.3 | -1287.0 | 43.7 | 2.66E-08 | -1713.7 | -1730.5 | 16.8 | 4.90E-03 |
| **ACE best (A)** | **Total sites (171,301)** | | | | **Aging I (500)** | | | **Aging II (795)** | | | | |
| **Fixed effects** | **B** | **SE** | **t** | **p** | **B** | **SE** | **t** | **p** | **B** | **SE** | **t** | **p** |
| Intercept  (Open Seas) | 0.070 | 0.000 | 156.32 | 0.00E+00 | 0.223 | 0.014 | 16.11 | 2.22E-58 | 0.123 | 0.009 | 13.99 | 1.86E-44 |
| Island | -0.010 | 0.001 | -16.08 | 3.54E-58 | -0.101 | 0.020 | -5.02 | 5.13E-07 | -0.018 | 0.015 | -1.20 | 2.31E-01 |
| N. Shelf | -0.002 | 0.001 | -1.47 | 1.41E-01 | -0.027 | 0.035 | -0.77 | 4.42E-01 | 0.036 | 0.023 | 1.57 | 1.16E-01 |
| N. Shore | 0.000 | 0.001 | 0.10 | 9.21E-01 | -0.042 | 0.020 | -2.06 | 3.94E-02 | 0.041 | 0.016 | 2.61 | 8.96E-03 |
| S. Shelf | -0.003 | 0.001 | -2.46 | 1.38E-02 | -0.046 | 0.041 | -1.13 | 2.60E-01 | -0.019 | 0.024 | -0.79 | 4.31E-01 |
| S. Shore | 0.000 | 0.001 | -0.12 | 9.03E-01 | -0.013 | 0.022 | -0.59 | 5.56E-01 | 0.040 | 0.017 | 2.33 | 1.96E-02 |
| **Random Effects** | **Intercept** | **Residual** | **ρ** | **--** | **Intercept** | **Residual** | **ρ** | **--** | **Intercept** | **Residual** | **ρ** | **--** |
| Variance | 0.008 | 0.004 | 0.653 | -- | 0.022 | 0.007 | 0.758 | -- | 0.020 | 0.005 | 0.796 | -- |
| **Model compare** | **Equal** | **Full** | **Δ χ2 (5)** | **p** | **Equal** | **Full** | **Δ χ2 (5)** | **p** | **Equal** | **Full** | **Δ χ2 (5)** | **p** |
| Deviance | -615899.2 | -616238.7 | 339.5 | 3.19E-71 | -1113.1 | -1141.0 | 27.9 | 3.81E-05 | -2088.4 | -2107.8 | 19.4 | 1.62E-03 |

*Note.* Models fitted in lme (version 1.1-21; Bates, Mächler, Bolker, & Walker, 2015); Aging I ,1217 CpG sites (717 ADE best, 500 ACE best) from Wang et al. (Wang et al., 2018); Aging II, 1934 CpG sites (1139 ADE best, 795 ACE best) from Tan et al. (Tan et al., 2016). Equal = model equating estimates across location ; Full = model freeing estimates across location.

*p* ≤ 4.52E-02

**Table S2. Multi-level regression results of heritability and shared environmental effects across age 69 and 79 years (“best” model estimates).**

|  |  |  | Aging 1 vs Background | | | | Aging II vs Background | | | | Clock sites vs Background | | | | Low Stability vs Background | | | |
| --- | --- | --- | --- | --- | --- | --- | --- | --- | --- | --- | --- | --- | --- | --- | --- | --- | --- | --- |
|  |  |  | B | se | Δχ^2^ (1) | p | B | se | Δχ^2^ (1) | p | B | se | Δχ^2^ (1) | p | B | se | Δχ^2^ (1) | p |
| Variance Components | A+D (ADE) | B_0_ | 0.237 | 0.000 | --- | --- | 0.237 | 0.000 | --- | --- | 0.237 | 0.000 | --- | --- | 0.236 | 0.000 |  |  |
|  |  | B_1_ | 0.157 | 0.006 | 735.6 | 5.43E-162 | 0.084 | 0.005 | 330.7 | 6.77E-74 | 0.050 | 0.006 | 75.2 | 4.25E-18 | 0.101 | 0.004 | 690.1 | 4.25E-152 |
|  | A (ACE) | B_0_ | 0.065 | 0.000 | --- | --- | 0.066 | 0.000 |  |  | 0.066 | 0.000 |  |  | 0.066 | 0.000 |  |  |
|  |  | B_1_ | 0.119 | 0.005 | 662.7 | 3.87E-146 | 0.067 | 0.004 | 335.3 | 6.74E-75 | 0.030 | 0.005 | 39.5 | 3.28E-10 | 0.137 | 0.005 | 676.4 | 4.05E-149 |
|  | C (ACE) | B_0_ | 0.102 | 0.000 | --- | --- | 0.102 | 0.000 |  |  | 0.102 | 0.000 |  |  | 0.102 | 0.000 |  |  |
|  |  | B_1_ | 0.046 | 0.003 | 177.3 | 1.88E-40 | 0.032 | 0.003 | 136.5 | 1.55E-31 | 0.022 | 0.004 | 37.6 | 8.68E-10 | 0.030 | 0.004 | 58.5 | 2.03E-14 |
| Absolute Variances | A+D (ADE) | B_0_ | 0.081 | 0.000 | --- | --- | 0.081 | 0.000 |  |  | 0.081 | 0.000 |  |  | 0.081 | 0.000 |  |  |
|  |  | B_1_ | 0.010 | 0.005 | 3.8 | 5.13E-02 | 0.025 | 0.004 | 39.6 | 3.12E-10 | 0.034 | 0.005 | 48.0 | 4.26E-12 | -0.019 | 0.003 | 34.0 | 5.51E-09 |
|  | E (ADE) | B_0_ | 0.205 | 0.000 | --- | --- | 0.205 | 0.000 |  |  | 0.204 | 0.000 |  |  | 0.205 | 0.000 |  |  |
|  |  | B_1_ | -0.077 | 0.007 | 141.2 | 1.45E-32 | -0.008 | 0.005 | 2.1 | 1.47E-01 | 0.036 | 0.006 | 30.2 | 3.90E-08 | -0.073 | 0.004 | 283.5 | 1.30E-63 |
|  | A (ACE) | B_0_ | 0.020 | 0.000 | --- | --- | 0.020 | 0.000 |  |  | 0.020 | 0.000 |  |  | 0.020 | 0.000 |  |  |
|  |  | B_1_ | 0.021 | 0.003 | 41.6 | 1.12E-10 | 0.020 | 0.003 | 62.9 | 2.17E-15 | 0.014 | 0.003 | 16.9 | 3.94E-05 | 0.010 | 0.004 | 7.9 | 4.94E-03 |
|  | C (ACE) | B_0_ | 0.024 | 0.000 | --- | --- | 0.024 | 0.000 |  |  | 0.024 | 0.000 |  |  | 0.024 | 0.000 |  |  |
|  |  | B_1_ | 0.008 | 0.002 | 13.8 | 2.03E-04 | 0.011 | 0.002 | 46.3 | 1.01E-11 | 0.016 | 0.002 | 54.6 | 1.48E-13 | -0.004 | 0.002 | 3.2 | 7.36E-02 |
|  | E (ACE) | B_0_ | 0.172 | 0.000 | --- | --- | 0.172 | 0.000 |  |  | 0.172 | 0.000 |  |  | 0.172 | 0.000 |  |  |
|  |  | B_1_ | -0.037 | 0.007 | 27.8 | 1.35E-07 | 0.009 | 0.006 | 2.6 | 1.07E-01 | 0.064 | 0.007 | 78.8 | 6.87E-19 | -0.069 | 0.008 | 75.1 | 4.47E-18 |

*Note.* Unconditional intercept-only models fitted in *lme* (version 1.1-21; Bates et al., 2015); B_0_ = intercept (background CpGs not in the Aging/Clock/Low stability set); B_1_ = difference in components/absolute variances in the Aging/Clock set vs background CpGs; ADE best included 187,535 total sites, ADE best included 171,301 total sites; Aging I ,1217 CpG sites (717 ADE best, 500 ACE best) from Wang et al. (Wang et al., 2018); Aging II, 1934 CpG sites (1139 ADE best, 795 ACE best) from Tan et al. (Tan et al., 2016); Clock, 1190 unique CpG sites (728 ADE best, 462 ACE best) from the Hannum (Hannum et al., 2013), Horvath (Horvath, 2013), Levine (Levine et al., 2018), or Zhang clocks (Zhang et al., 2019); Low Stability comparison included 2020 sites (1638 ADE best, 382 ACE best).

*p* ≤ 4.52E-02

**Table S3. Multi-level regression results of familial effects by clock type across age 69 and 79 years.**

| **Fixed effects** | **Variance Components** | | | | **Absolute Variances** | | | |
| --- | --- | --- | --- | --- | --- | --- | --- | --- |
| **A+D (ADE)** | **B** | **SE** | **t** | **p** | **B** | **SE** | **t** | **p** |
| B_0_ (Zhang) | 0.333 | 0.010 | 33.58 | 3.13E-247 | 0.140 | 0.009 | 14.91 | 2.79E-50 |
| B Levine | -0.071 | 0.014 | -5.07 | 4.00E-07 | -0.039 | 0.013 | -2.95 | 3.18E-03 |
| B Horvath | -0.080 | 0.016 | -5.02 | 5.16E-07 | -0.040 | 0.015 | -2.67 | 7.54E-03 |
| B Hannum | -0.018 | 0.040 | -0.45 | 6.53E-01 | -0.030 | 0.038 | -0.79 | 4.31E-01 |
| **Random Effects** | **B_0_** | **Resid** | **ρ** | **−−** | **B_0_** | **Resid** | **ρ** | **−−** |
| Variances | 0.019 | 0.015 | 0.559 | **−−** | 0.022 | 0.003 | 0.879 | **−−** |
| **Model compare** | **Equal** | **Full** | **Δ χ2 (3)** | **p** | **Equal** | **Full** | **Δ χ2 (3)** | **p** |
| Deviance | -1045.0 | -1079.9 | 34.9 | 1.28E-07 | -2305.6 | -2316.6 | 11.0 | 1.17E-02 |
| **E (ADE)** |  |  |  |  | **B** | **SE** | **t** | **p** |
| B_0_ (Zhang) | **−−** | **−−** | **−−** | **−−** | 0.239 | 0.011 | 21.70 | 2.11E-104 |
| B Levine | **−−** | **−−** | **−−** | **−−** | 0.002 | 0.015 | 0.12 | 9.07E-01 |
| B Horvath | **−−** | **−−** | **−−** | **−−** | -0.001 | 0.018 | -0.06 | 9.52E-01 |
| B Hannum | **−−** | **−−** | **−−** | **−−** | 0.015 | 0.045 | 0.33 | 7.39E-01 |
| **Random Effects** | **−−** | **−−** | **−−** | **−−** | **B_0_** | **Resid** | **ρ** | **--** |
| Variance | **−−** | **−−** | **−−** | **−−** | 0.028 | 0.008 | 0.776 | **--** |
| **Model compare** | **−−** | **−−** | **−−** | **−−** | **Equal** | **Full** | **Δ χ2 (3)** | **p** |
| Deviance | **−−** | **−−** | **−−** | **−−** | -1358.5 | -1358.6 | 0.1 | 9.92E-01 |
| **A (ACE)** | **B** | **SE** | **t** | **p** | **B** | **SE** | **t** | **p** |
| B_0_ (Zhang) | 0.116 | 0.009 | 13.03 | 8.55E-39 | 0.038 | 0.005 | 7.95 | 1.88E-15 |
| B Levine | -0.041 | 0.013 | -3.07 | 2.15E-03 | -0.010 | 0.007 | -1.35 | 1.76E-01 |
| B Horvath | -0.029 | 0.015 | -1.94 | 5.26E-02 | -0.008 | 0.008 | -1.02 | 3.08E-01 |
| B Hannum | 0.000 | 0.035 | -0.01 | 9.91E-01 | 0.022 | 0.019 | 1.17 | 2.41E-01 |
| **Random Effects** | **B_0_** | **Resid** | **ρ** | **−−** | **B_0_** | **Resid** | **ρ** | **−−** |
| Variance | 0.012 | 0.006 | 0.680 | **−−** | 0.004 | 0.001 | 0.779 | **−−** |
| **Model compare** | **Equal** | **Full** | **Δ χ2 (3)** | **p** | **Equal** | **Full** | **Δ χ2 (3)** | **p** |
| Deviance | -1376.1 | -1386.4 | 10.3 | 1.62E-02 | -2723.3 | -2727.6 | 4.3 | 2.31E-01 |
| **C (ACE)** | **B** | **SE** | **t** | **p** | **B** | **SE** | **t** | **p** |
| B_0_ (Zhang) | 0.157 | 0.006 | 25.00 | 6.70E-138 | 0.050 | 0.004 | 14.26 | 3.70E-46 |
| B Levine | -0.059 | 0.009 | -6.28 | 3.32E-10 | -0.022 | 0.005 | -4.22 | 2.45E-05 |
| B Horvath | -0.059 | 0.011 | -5.60 | 2.19E-08 | -0.017 | 0.006 | -2.82 | 4.75E-03 |
| B Hannum | 0.018 | 0.025 | 0.72 | 4.71E-01 | 0.006 | 0.014 | 0.46 | 6.44E-01 |
| **Random Effects** | **B_0_** | **Resid** | **ρ** | **--** | **B_0_** | **Resid** | **ρ** | **−−** |
| Variance | 0.005 | 0.005 | 0.481 | -- | 0.002 | 0.001 | 0.657 | **−−** |
| **Model compare** | **Equal** | **Full** | **Δ χ2 (3)** | **p** | **Equal** | **Full** | **Δ χ2 (3)** | **p** |
| Deviance | -1699.6 1699.60 | -1751.8 | 52.2 | 2.72E-11 | -3045.6 | -3066.2 | 20.6 | 1.27E-04 |
| **E (ACE)** |  |  |  |  | **B** | **SE** | **t** | **p** |
| B_0_ (Zhang) | **−−** | **−−** | **−−** | **−−** | 0.224 | 0.013 | 16.60 | 7.14E-62 |
| B Levine | **−−** | **−−** | **−−** | **−−** | 0.008 | 0.020 | 0.40 | 6.87E-01 |
| B Horvath | **−−** | **−−** | **−−** | **−−** | 0.033 | 0.023 | 1.45 | 1.46E-01 |
| B Hannum | **−−** | **−−** | **−−** | **−−** | 0.097 | 0.053 | 1.83 | 6.68E-02 |
| **Random Effects** | **−−** | **−−** | **−−** | **−−** | **B_0_** | **Resid** | **ρ** | **−−** |
| Variance | **−−** | **−−** | **−−** | **−−** | 0.031 | 0.006 | 0.838 | **−−** |
| **Model compare** | **−−** | **−−** | **−−** | **−−** | **Equal** | **Full** | **Δ χ2 (3)** | **p** |
| Deviance | **−−** | **−−** | **−−** | **−−** | -974.2 | -979.1 | 4.9 | 1.79E-01 |

*Note*. Models fitted in lme (version 1.1-21; Bates et al., 2015); 1190 unique CpG sites with 59 of 71 sites available from the Hannum clock (Hannum et al., 2013), 312 of 353 sites from the Horvath clock (Horvath, 2013), and 443 of 513 sites from the Levine clock(Levine et al., 2018), and 455 of 514 sites from the Zhang clock (Zhang et al., 2019). Resid = residual. Equal = model equating estimates across clock type; Full = model freeing estimates across clock type.

**Figure S1. Change in variance for 2020 CpGs selected for low stability (1638 ADE best, 382 ACE best).**

|  | **Variance Components** | **Absolute Variances** |
| --- | --- | --- |
| **ADE best** | 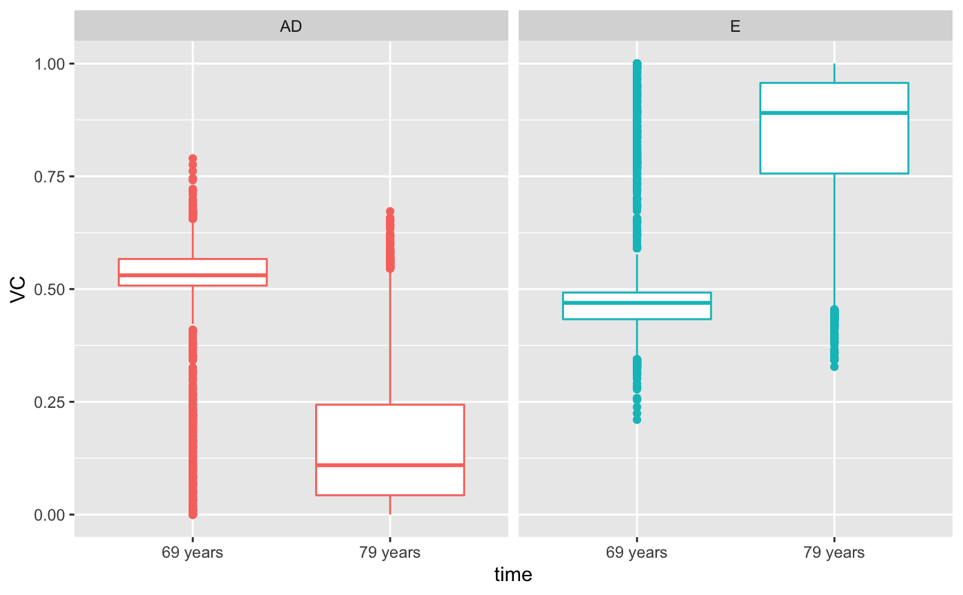 | 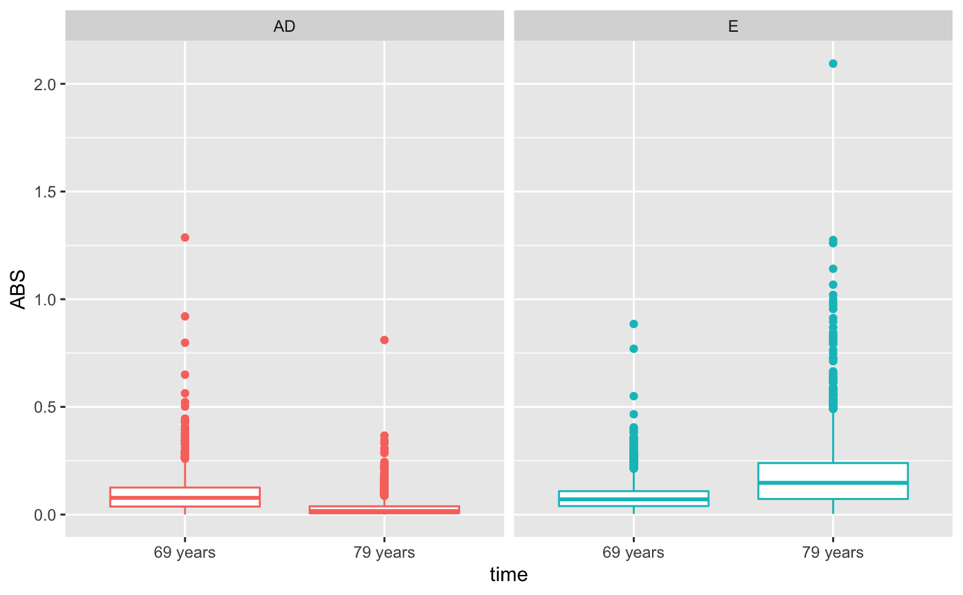 |
| **ACE best** | 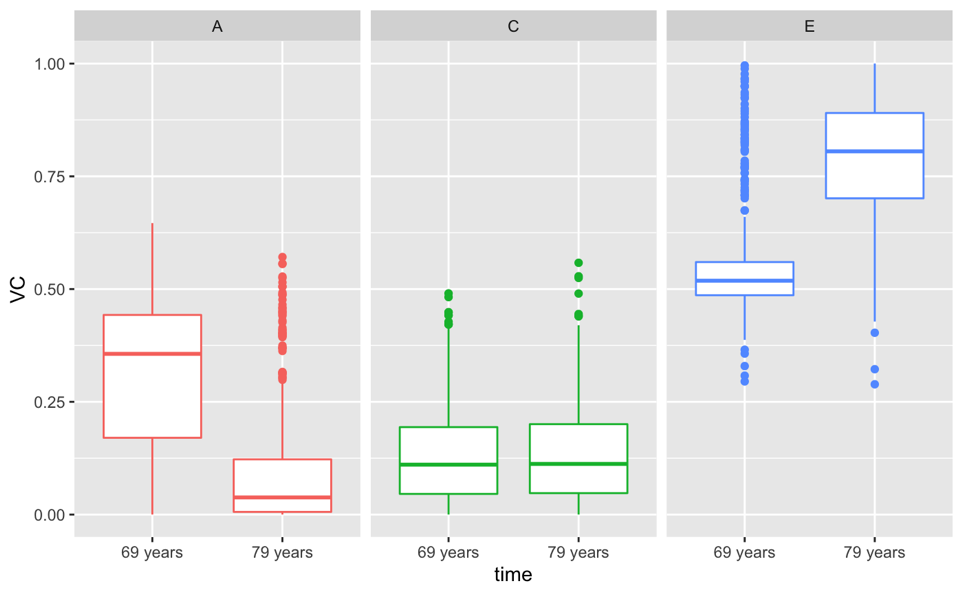 | 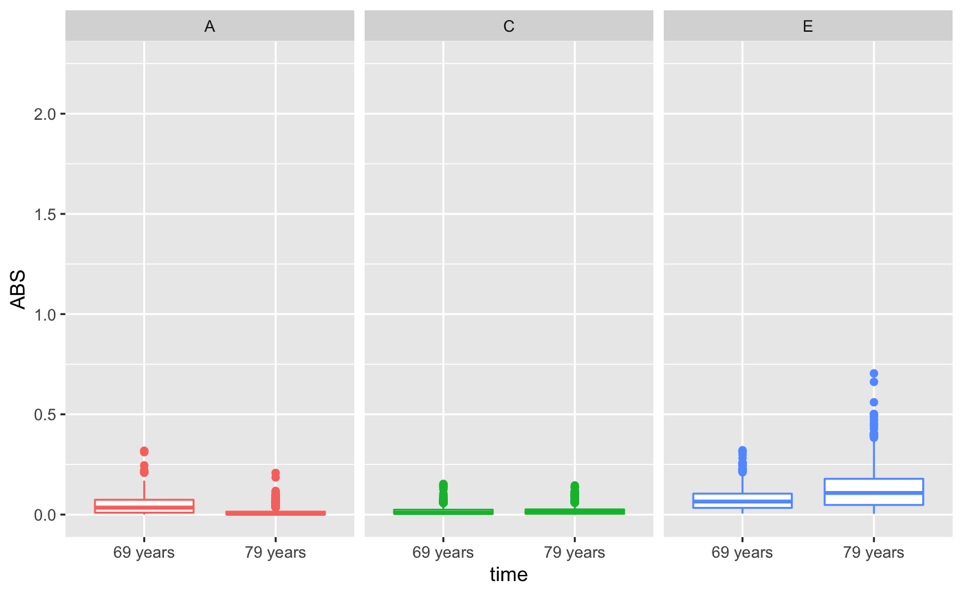 |

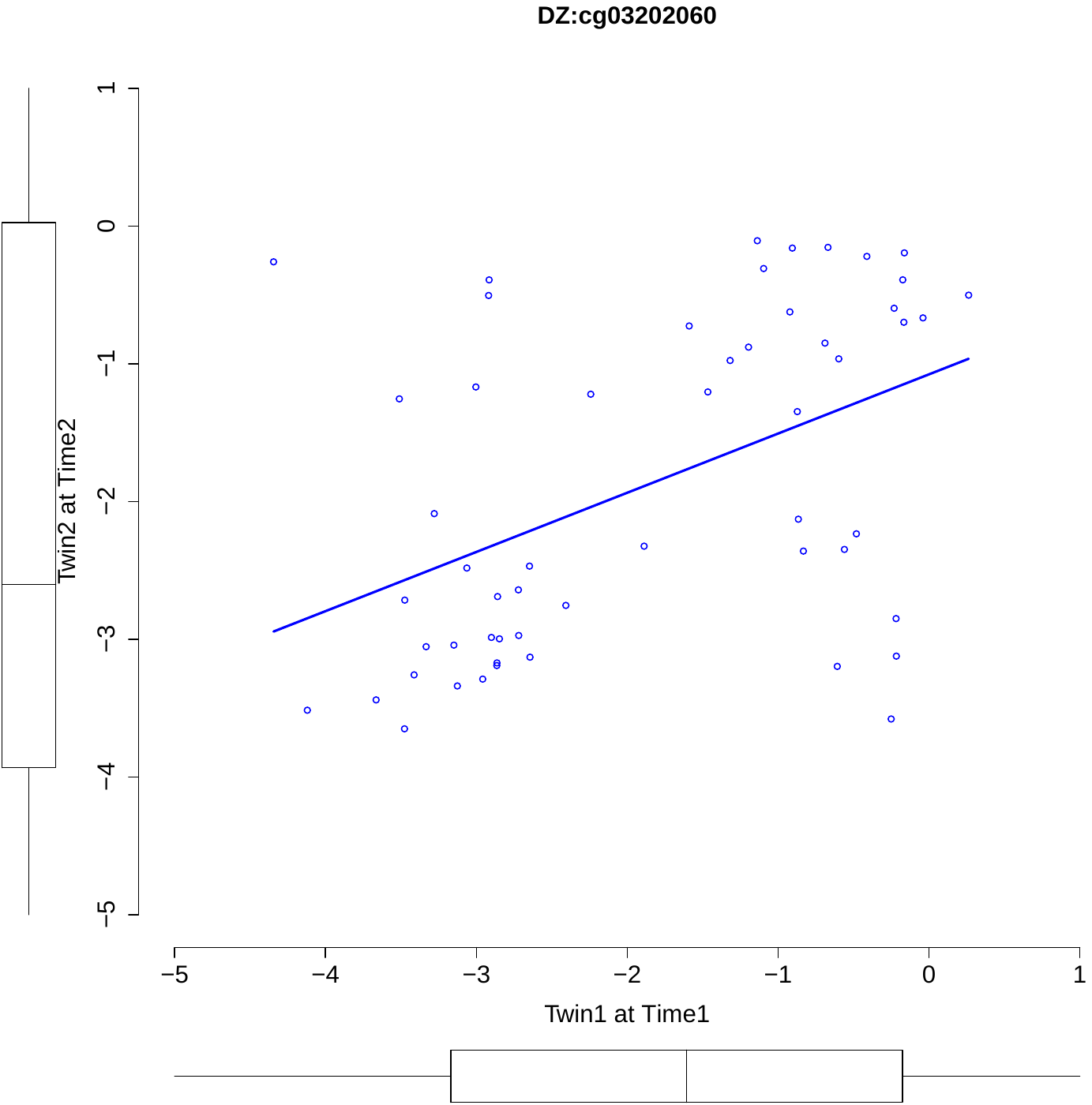
**Figure S2. Scatterplot of M-values across time by twins (MZ, DZ) for cg03202060 (panels a and b) and cg07677296 (panels c and d).**

*Note***.** Twin1 at time 1 on *x*-axis, and twin 2 at time 2 on *y-*axis.

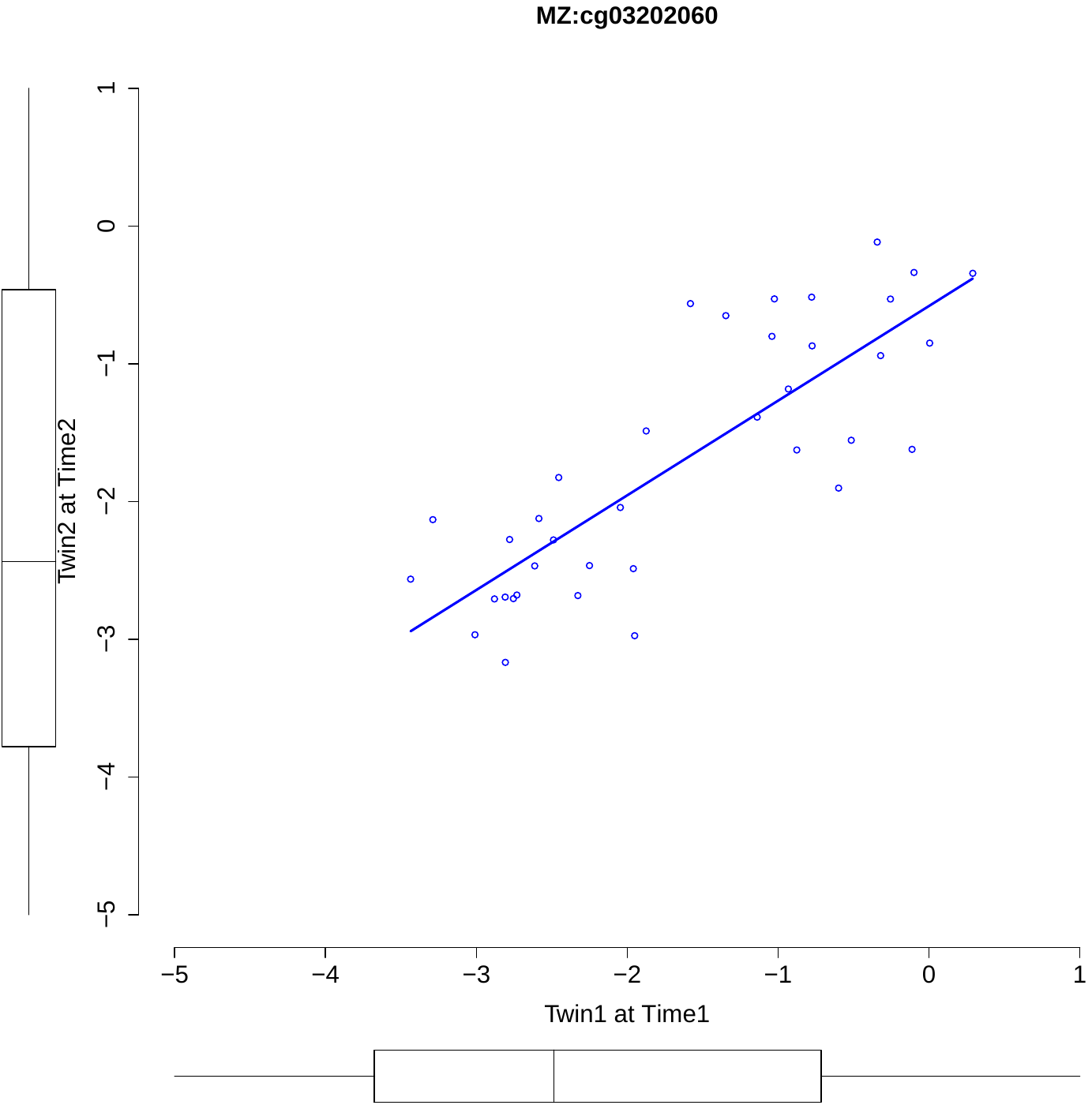

**b**

**a**

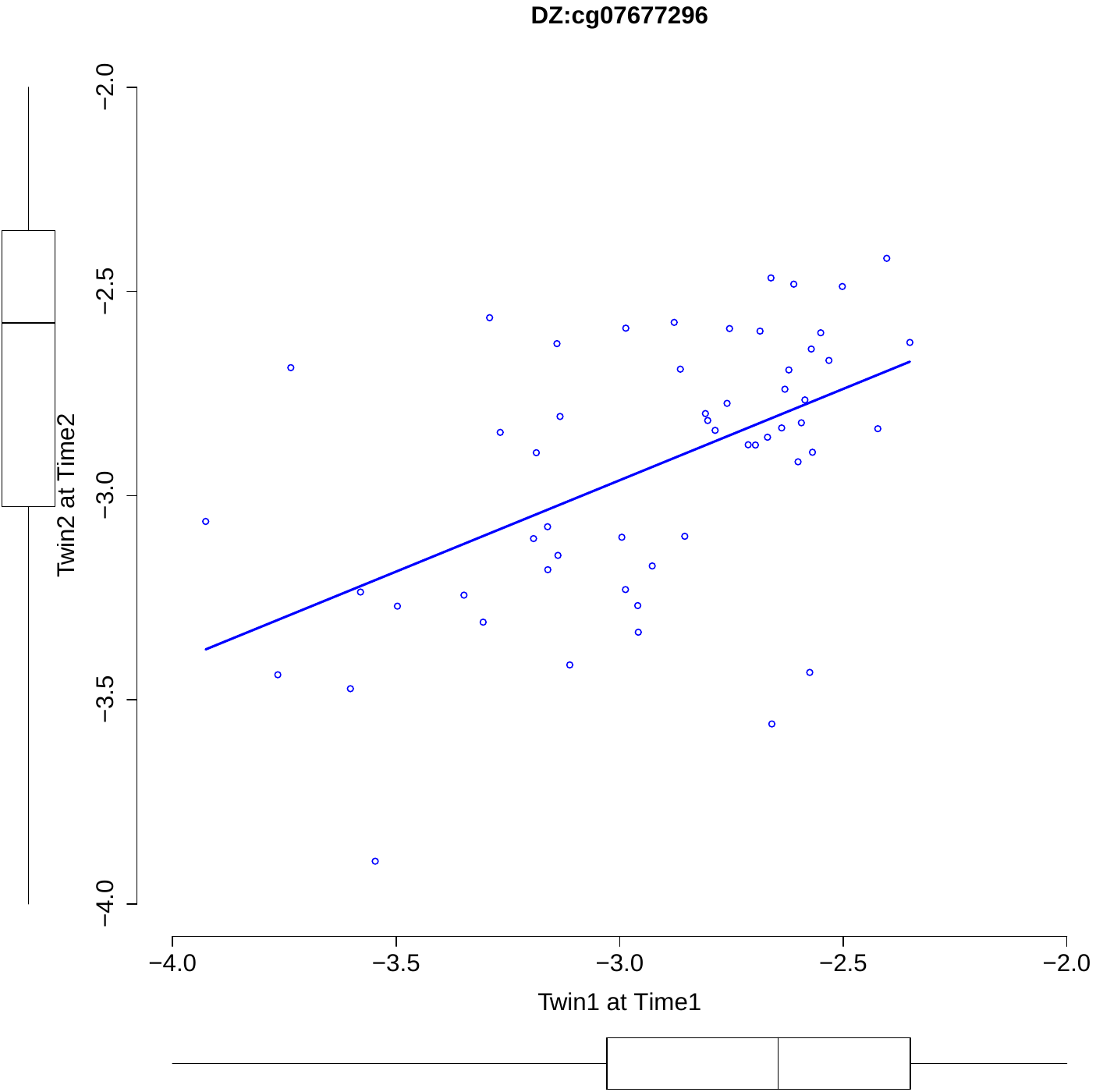

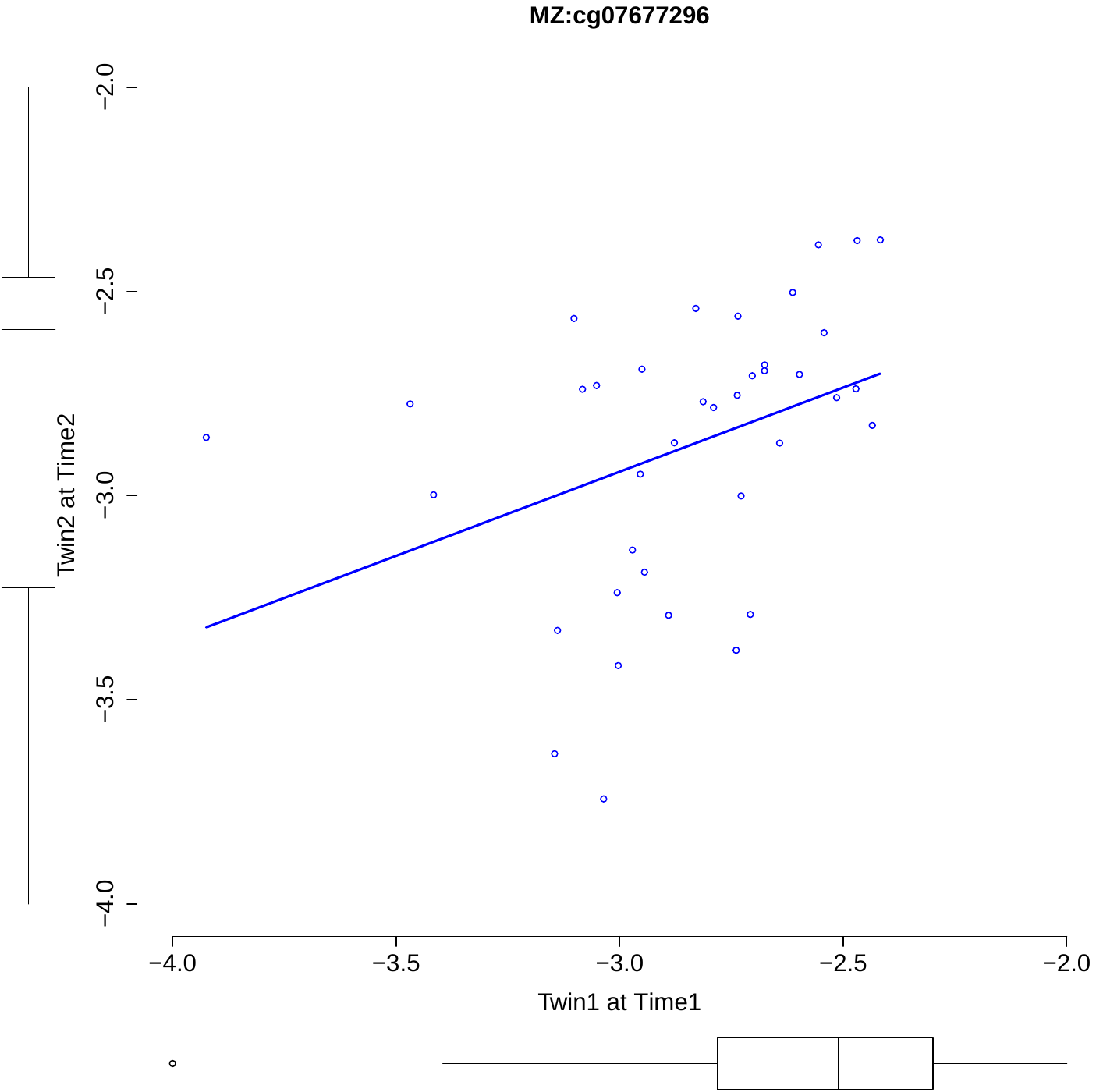

**c**

**d**
